# Genome-wide expression QTL mapping reveals the highly dynamic regulatory landscape of a major wheat pathogen

**DOI:** 10.1101/2023.07.14.549109

**Authors:** Leen Nanchira Abraham, Daniel Croll

## Abstract

**Background:** In agricultural ecosystems, outbreaks of diseases are frequent and pose a significant threat to food security. A successful pathogen undergoes a complex and well-timed sequence of regulatory changes to avoid detection by the host immune system, hence well-tuned gene regulation is essential for survival. However, the extent to which the regulatory polymorphisms in a pathogen population provide an adaptive advantage is poorly understood.

**Results:** We used *Zymoseptoria tritici*, one of the most important pathogens of wheat, to generate a genome-wide map of regulatory polymorphism governing gene expression. We investigated genome-wide transcription levels of 146 strains grown under nutrient starvation and performed expression quantitative trait loci (eQTL) mapping. We identified *cis*-eQTLs for 65.3% of all genes and the majority of all eQTL loci are within 2kb upstream and downstream of the transcription start site (TSS). We also show that polymorphism in different gene elements contributes disproportionally to gene expression variation. Investigating regulatory polymorphism in gene categories, we found an enrichment of regulatory variants for genes predicted to be important for fungal pathogenesis but with comparatively low effect size, suggesting a separate layer of gene regulation involving epigenetics. We also show that previously reported trait-associated SNPs in pathogen populations are frequently *cis*-regulatory variants of neighboring genes with implications for the trait architecture.

**Conclusions:** Overall, our study provides extensive evidence that single populations segregate large-scale regulatory variation and are likely to fuel rapid adaptation to resistant hosts and environmental change.

## Background

The control of gene expression is essential for the development and survival of organisms. The foundation of gene regulation is the interaction of regulatory proteins with specific DNA (regulatory) sequences in the coding or non-coding regions of the genome. Transcription factors (TFs) modulate gene expression by binding to transcription factor binding sites in regulatory regions [1, 2]. For instance, TF binding to the promoter near the transcription start site (TSS) helps to initiate transcription by forming a transcription initiation complex [3]. The regulatory region of a gene was thought to be usually located upstream of the TSS. However, comprehensive studies across eukaryotes showed that both the 5’ and 3’-untranslated regions, introns, and even coding regions can act as regulatory sequences [4]. In addition to regulatory sequences, eukaryotic gene regulation is also governed by chromatin structure and histones [5].

Even though core mechanisms of gene expression are conserved across eukaryotes, there is substantial variation in gene expression within species. Individual genotypes can carry adaptive regulatory mutations adjusting gene expression to different environmental cues. In clinical strains of yeasts, promoter variants can cause the upregulation of biofilm suppressor genes and increases pathogenicity [6]. Different consumption rates of aspartic and glutamic acid during the fermentation of yeast hybrids are mediated by the reduced binding affinity of the transcriptional activator protein (Uga3) underpinned by single nucleotide polymorphisms (SNPs) in the binding region [7]. A synonymous mutation can increase the fitness of *Pseudomonas fluorescens* by increasing gene expression [8]. Hence, understanding how regulatory variation can promote adaptive evolution is important and requires population-scale approaches. However, most evidence relies on a few model organisms such as yeasts [9–11], *Drosophila* [12, 13], humans [14–16] or *C. elegans* [17] and we lack comprehensive studies of regulatory variation in many ecologically relevant species.

Mapping of regulatory mutations in model organisms generated fundamental insights into the extent of regulatory polymorphism and gene regulatory networks. SNPs, insertions, deletions, copy-number variants, as well as transposable elements (TE) can act as a genetic variant [9, 18, 19]. Genetic variants associated with mRNA level variation are called expression quantitative trait loci (eQTLs). eQTLs can be located in regulatory sequences, such as promoters, enhancers, and splice sites, or gene elements such as 5 or 3’ UTR, exons, and introns. Most eQTLs are acting in *cis* on the neighboring genes. But eQTLs can also be in genes typically encoding regulatory proteins that interact with *cis*-regulatory sequences on the same or different chromosome (*i.e. trans* eQTLs) (Kita, Venkataram, Zhou, & Fraser, 2017; Lutz et al., 2019; Williams et al., 2007). Species differ in their organization of regulatory regions. For instance, in yeasts, regulatory variants are enriched upstream of the TSS (i.e., the promoter) and 3’UTR whereas in human and *Drosophila* SNPs located near the TSS and in the 5′UTRs are more likely to be eQTLs [9, 20]. *Cis*-eQTLs tend to be more common and have larger effect sizes on gene expression variation than *trans-*eQTLs. Exceptions include for example the plant pathogenic fungi *Botrytis cinerea* and *Coprinopsis cinerea* [21–23]. Plant pathogens often harbor also polymorphism for gene expression during pathogenesis with likely strong selective advantages [24–27].

Plant pathogens cope with multiple environmental stressors over the life cycle with well-timed sequences of regulatory changes. During plant infection, pathogens tightly control gene expression to circumvent recognition by the host [28–31]. Pathogens secrete effectors (*i.e.* small secreted proteins) at the onset of host infection serving to manipulate the host cell biology, repress host immune responses, or shield the pathogen to support growth and colonization [32–35]. Because of their prime role during infection, effector gene expression is highly upregulated upon contact with the host [28]. Carbohydrate-active enzymes (CAZyme) are a second major component of fungal pathogenicity and serve to break down plant cell wall components to aid host colonization. *Zymoseptoria tritici*, a major pathogen of wheat shows evidence for intra-specific regulatory variation and high genetic diversity from single field populations to continents [36, 37]. Populations show a rapid decay in linkage disequilibrium which helps increase the power of mapping approaches and narrows down associations [38–40]. The insertion of a TE led to the downregulation of a recognized effector gene and enabled the strain to evade recognition by the host [41]. Similarly, differences in the production of melanin were shown to be controlled by epigenetic regulation of nearby TEs underlining the significance of regulatory variants associated with gene expression variation within pathogen populations [42, 43] TE-associated polymorphisms and effects on gene expression are widespread in the genome and underpin phenotypic trait variation even within single field populations [44, 45]

In this study, we map regulatory variation at the genome-wide scale in the fungal pathogen of wheat *Z. tritici*. We associate variation in gene transcription levels with genetic variation from strains collected from a highly diverse field population. We assess *cis* and *trans-*regulatory variation segregating for different gene functions and genomic locations. Furthermore, we test whether different functional gene categories important for pathogenesis show differences in the level of segregating regulatory variation. Finally, we analyze contributions of regulatory variations on phenotypic trait variation within the species.

## Results

### Polymorphism analyses of the mapping population

We analyzed 146 *Z. tritici* isolates from a field population in Switzerland using whole-genome and transcriptome sequencing [36, 44]. Variant calling on whole-genome sequences of the mapping population identified 543,046 SNPs and 39,356 indels after quality filtering. The isolates were highly diverse compared to the worldwide diversity of the pathogen (Figure 1A). We identified a strong skew of minor allele frequencies towards rare variants consistent with a large, recombining population (Suppl. Figure S1). A principal component analysis of genetic variants showed no evidence for population substructure among the analyzed isolates with only 2% of the cumulative variance explained by PC1 (Figure 1B, Suppl. Figure S2). We quantified genome-wide transcription of the 146 *Z. tritici* isolates under nutrient starvation conditions. A PCA of gene expression showed minor clustering according to culturing and RNA collection batches and cumulative variance explained by PC1 was around 10% (Figure 1C, Suppl. Figure S3). The population showed high expression variation between different gene categories important for pathogenesis (Figure 1D). To account for heterogeneity in the mapping population, we included the first PC from the genetic substructure analysis, as well as PCs 1-10 from the gene expression analyses as covariates in the genome-wide mapping (Suppl. Fig. S3A-B). To correct for the minor batch effect, we included the RNA sequencing batch as a random effect in the association mapping analyses (Figure 1C).

**Figure 1.**
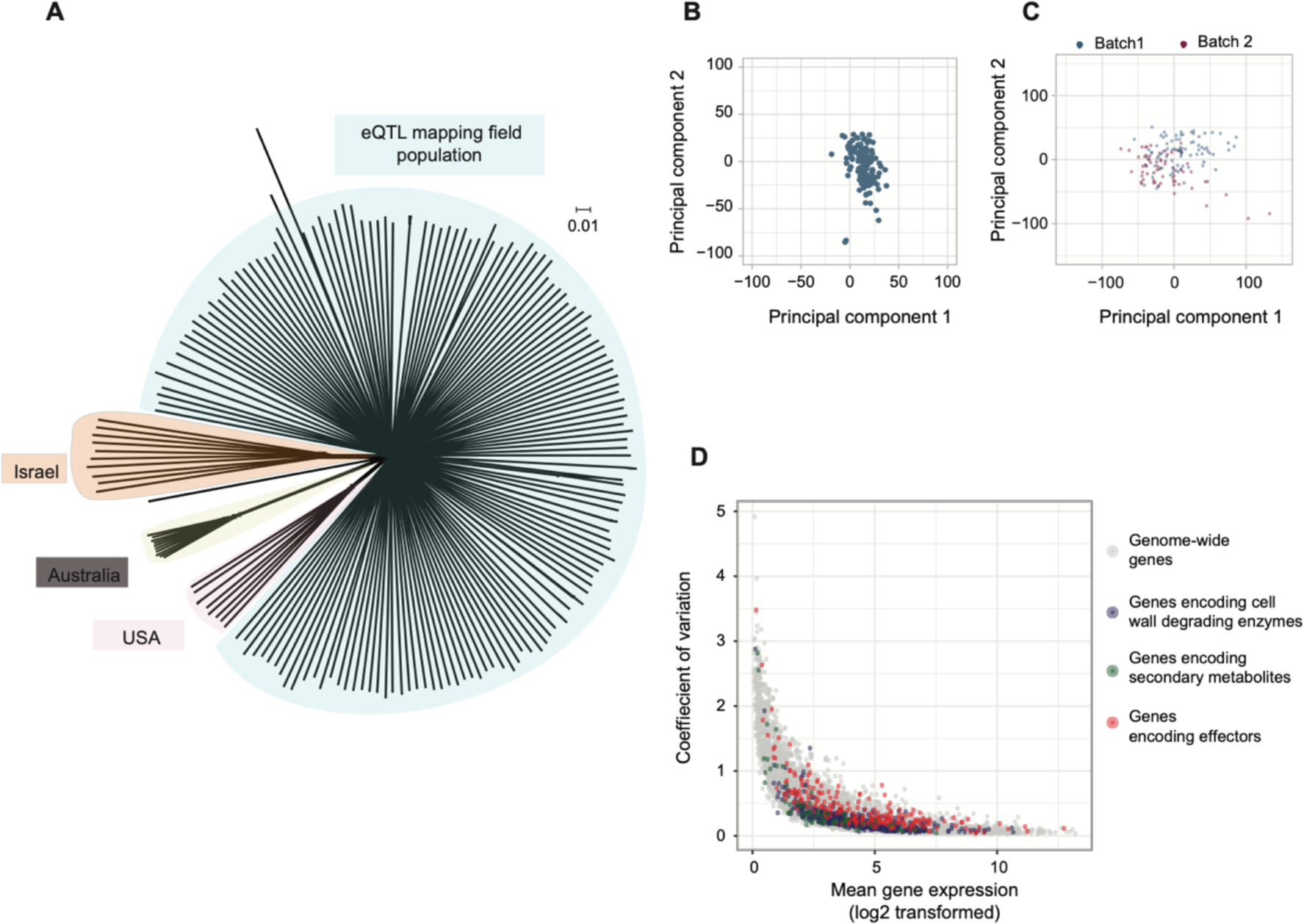
Genetic diversity of the mapping population. (A) Unrooted phylogenetic network of the mapping population and representative isolates from a global collection of *Z. tritici* populations. (B) Principal component analysis of genetic variation within the mapping population. (C) Principal component analysis of gene expression in the mapping population. Colors differentiate culture batches. (D) Coefficient of gene expression variation against the RPKM normalized values of gene expression (log2-transformed). Colors indicate different gene categories relevant for pathogenesis.

### Genome-wide regulatory variation

To identify *cis*-regulatory variation we associated the effect of polymorphisms around TSS to gene expression variation of the gene. The 10 kb *cis* window covers the majority of the gene length (mean gene length ∼1.6kb) and avoids significant overlap with neighboring genes (mean intergenic distance ∼1.7kb; Figure 2A). We observed that the majority of the significant *cis*-eQTLs mapped near the 10kb window around TSS even with an increase in cis window to 25kb from the TSS (Suppl. Figure S4A-B). We identified 11,083 *cis*-eQTLs (FDR >5%) across the genome with 65.3% (*n* = 7396) of the genes showing at least one *cis*-eQTL (Figure 2B, Suppl. Table S1). Among genes with a mapped *cis-*eQTL, 61.2% of the genes show a single *cis*-eQTL, a further 29.3%, 7.7% and 1.3% show two, three and four *cis-*eQTL, respectively (Suppl. Figure S5). The proportion of genes with at least one mapped eQTL was substantially higher on core chromosomes compared to accessory chromosomes (Figure 2C).

**Figure 2.**
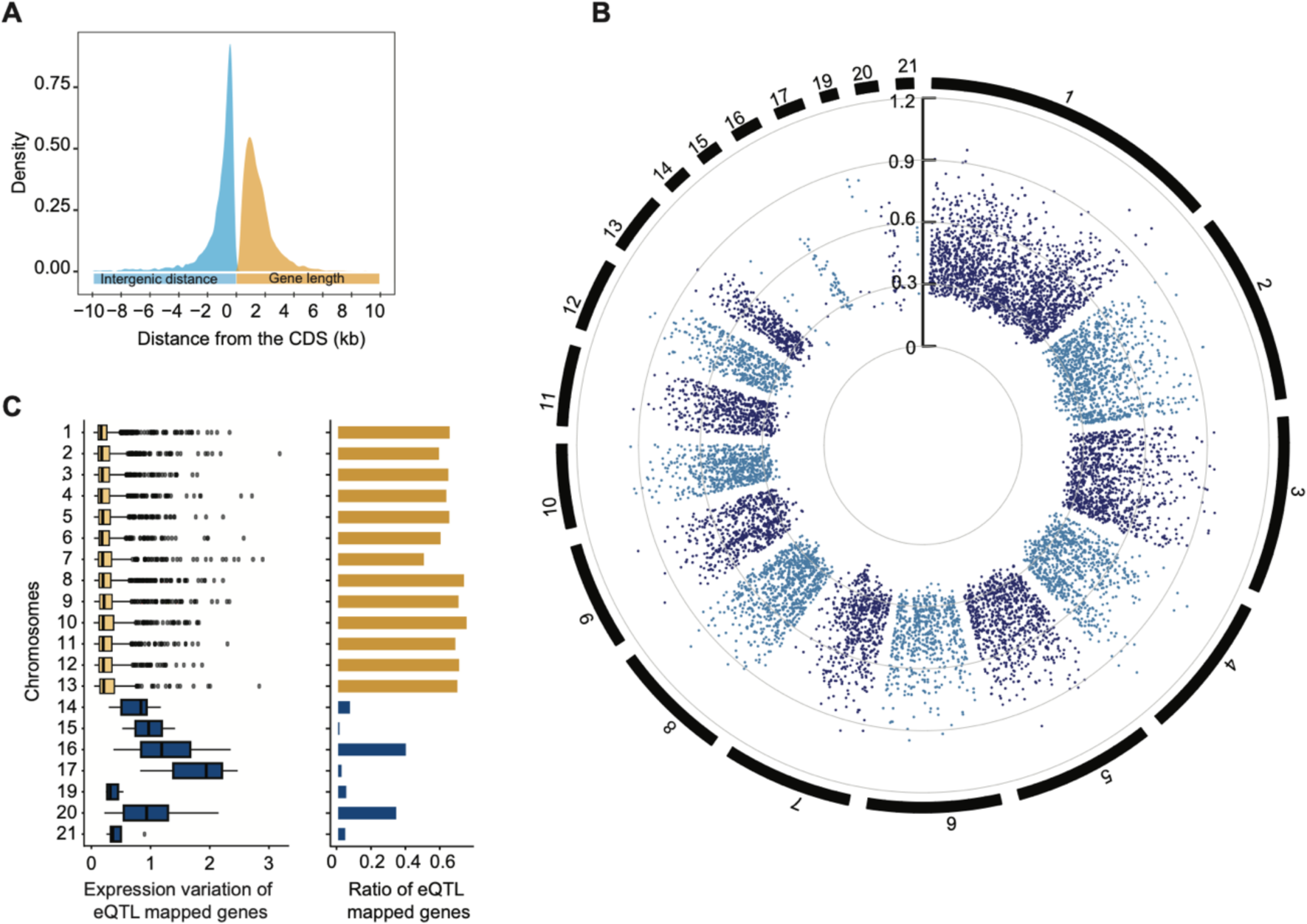
Genome-wide distribution of mapped *cis*-eQTLs. (A) Genome-wide distribution of gene length and intergenic distances. (B) Genome-wide distribution of mapped *cis*-eQTLs. The scale inside indicates the effect size of the eQTLs. (C) Distribution of expression variation and ratio of genes with an eQTL in core and accessory chromosomes. Gene expression on accessory chromosomes is normalized for chromosome presence-absence in the population.

### Contribution of gene features to regulatory variation

We analyzed whether *cis-*eQTL were enriched in specific gene elements. We observed varying densities of *cis*-eQTLs in the coding sequence, 3’UTR, 5’UTR, intron, upstream of TSS, and downstream of TES respectively (Figure 3A, 3B) with a higher density of *cis*-eQTLs in the coding sequence and an increase in density towards the 3’ end. To account for variation in SNP densities across the gene element, we assessed the enrichment of SNPs being mapped as an eQTL in different gene elements. We found an enrichment of *cis*-eQTL upstream of the TSS, 5’ UTR, and 3’UTR, regardless of the variant category (Figure 3C). In contrast to the enrichment pattern, the effect size is higher for *cis*-eQTLs mapped in exons and introns. Intron located indels mapped as *cis*-eQTLs showed a higher effect size than other *cis*-eQTLs (Figure 3D).

**Figure 3.**
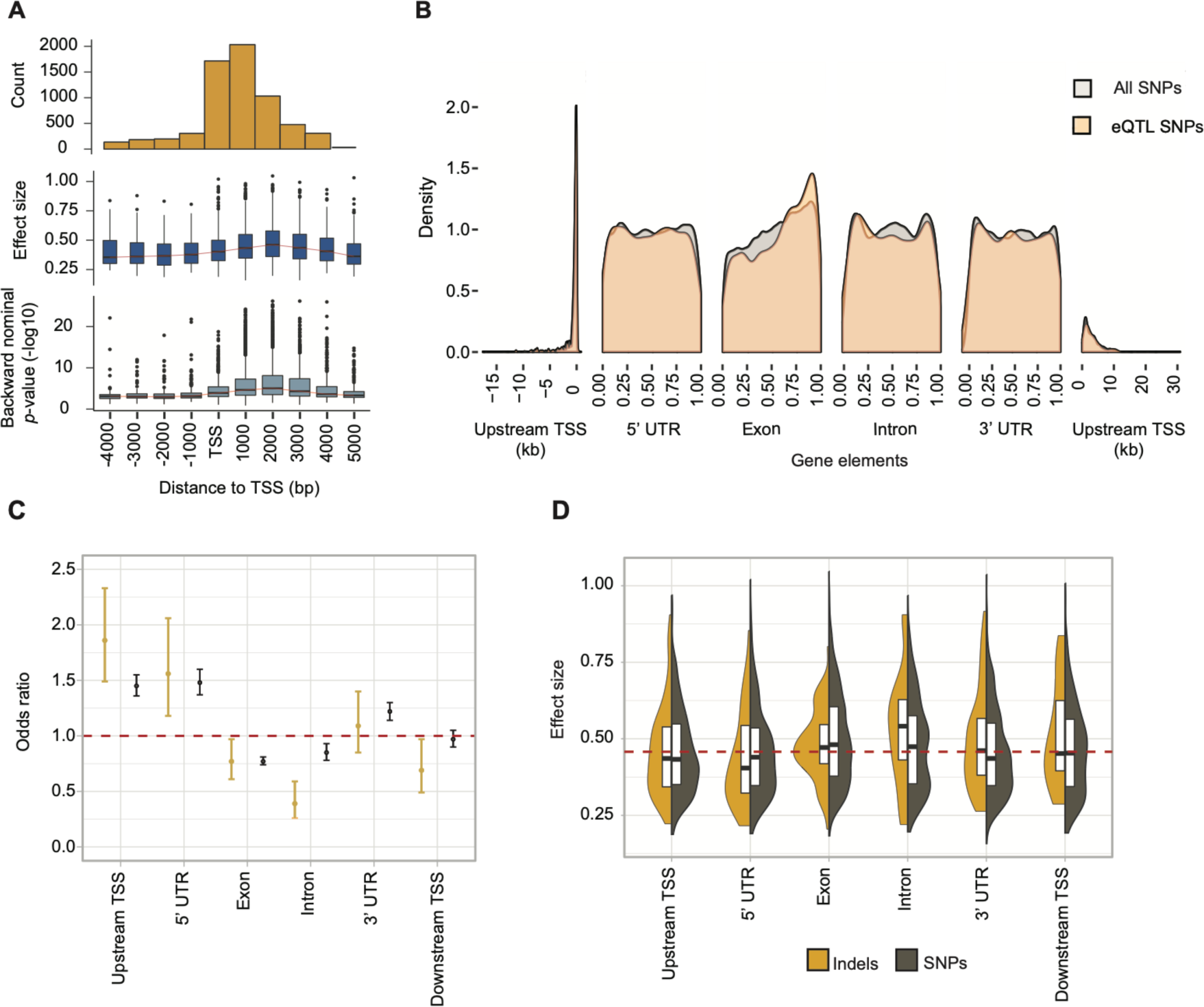
Localization of *cis*-eQTLs across gene elements. (A) Density of mapped eQTLs in *cis* windows. (B) Distribution of *cis*-eQTLs relative to different gene elements. 5’, UTR, exon and intron positions are shown scaled to 1. Upstream/downstream TSS positions are shown in kb. (C) Enrichment of *cis*-eQTLs in gene elements (odds ratio of observed over expected). Color distinguishes polymorphism types at the eQTL (yellow: indel; brown: SNP). (D) The effect size of high-impact eQTLs across gene elements.

Furthermore, we assessed SNPs mapped as *cis*-eQTLs between different gene categories encoding essential protein functions for plant pathogens to successfully infect and exploit host tissue. We selected gene categories encoding cell wall degrading enzymes (CAZymes), genes encoding predicted effectors, secondary metabolite gene clusters, major facilitator superfamily (MFS) genes, and genes highly upregulated during plant infection [29]. We analyzed the coefficient of gene expression variation (CV) across these gene categories and found that genes encoding effectors have higher CV than other gene categories whereas genes highly expressed *in planta* showed the least expression variation (Figure 4A). The percentage of genes with at least one mapped eQTL is lower for genes with high expression in *planta* (48%) followed by core biosynthetic genes of secondary metabolite gene clusters (54%), genes encoding candidate effectors (55%), genes encoding MFS transporters (62%) and genes encoding CAZymes (67%) (Figure 4B). Highly expressed genes *in planta* infection are more likely to have at least one *cis*-eQTL (95% confidence interval; Figure 4C) but with comparatively lower effect size (Figure 4D). We hypothesize that epigenetic regulation might be more prevalent than regulatory polymorphism in genes encoding effectors and highly expressed genes during plant infection. To test this, we investigated the coverage of different histone marks in the *cis* window of these gene categories in the reference genome. We observed a higher coverage of euchromatin histone marks H3K4m2 in the window of genes encoding effectors and genes reported with high expression i*n planta* compared to other gene categories (Figure 4E). H3K4m2 histone marks tend to be involved in the regulation of gene expression, which supports the hypothesis that gene categories mapped with low effect size regulatory *cis*-eQTLs are more likely regulated by epigenetics rather than *cis*-eQTL variants.

**Figure 4.**
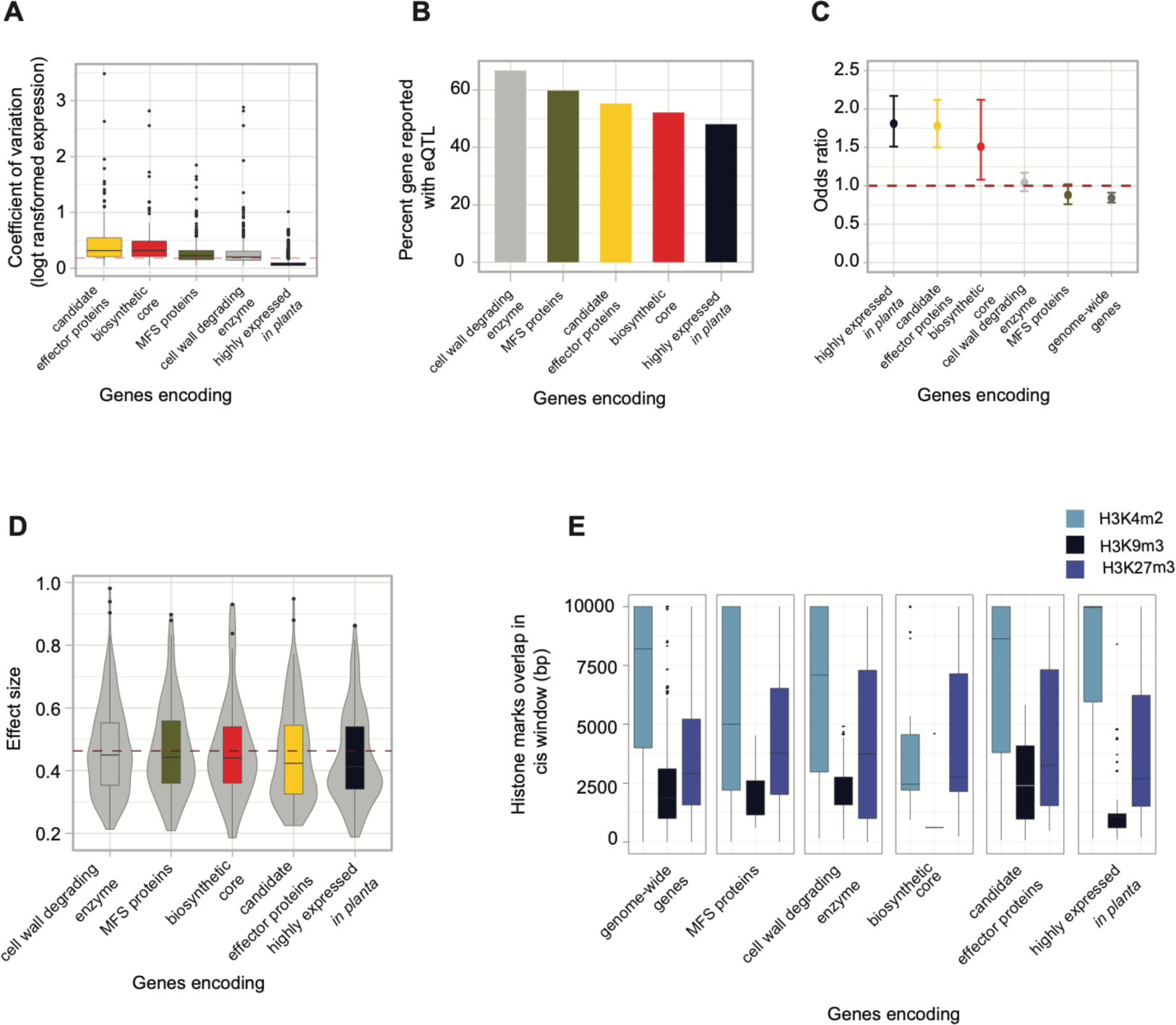
The proportion of cis-eQTLs identified across gene categories. (A) Expression variation of gene categories across gene categories. (B) Percentage of genes reported with at least one eQTL across gene categories. (C) Enrichment of cis-eQTLs across gene categories. (D) Effect size of high impact eQTLs across gene categories. The red line indicates the cis-eQTL effect size of genome-wide genes. (E) Overlap of three different histone methylation marks in cis windows of different gene categories.

### Networks of genes regulated by trans-eQTLs

We performed *trans-*eQTL mapping to identify regulatory variants associated with the expression of distal genes. The full permutation pass with stringent criteria identified 20 genes regulated by *trans-* eQTL with high confidence whereas the approximation pass with less multiple testing burden on *trans-* eQTLs reported 843 genes with at least one *trans-*eQTL (Suppl. Tables S2 and S3). The *trans-*eQTLs from the approximate pass were used to reconstruct genome-wide pattern of *trans-*eQTL occurrence. We found that polymorphisms on core chromosomes are more likely to influence genes on core chromosomes than accessory chromosomes. Whereas the proportion of genes mapped with at least a *trans-*eQTL does not differ between core and accessory chromosomes (Figure 5A)., in contrast to the findings for *cis*-eQTL occurrences. We also observed multiple *trans*-eQTLs from different chromosomes regulating a single gene implying a complex network of gene regulation spanning different chromosomes (Suppl. Table S3). An interesting locus reported as a *trans*-eQTL is a polymorphism in an exon of the gene 10_00067 located on chromosome 10 associated to the expression of the gene 21_00018 on chromosome 21 (Figure 5B). Both genes are predicted to be encoding alpha-tubulin suggesting coregulation of alpha-tubulin genes in the genome. A small proportion of *trans-* eQTLs (83 of 843) were found to be overlapping with *cis*-eQTLs. The effect size of eQTLs mapped with *cis* and *trans* effects is higher than the effect size of eQTLs having only a *cis* or *trans* effect (Figure 5C). These overlapping pairs are possibly explained by functional links between the *cis* and *trans* linked genes. Among overlapping *cis/trans* eQTLs, 46 are regulating the expression of *cis* and *trans* genes in the same orientation and 42 in opposite orientations (Figure 5D).

**Figure 5.**
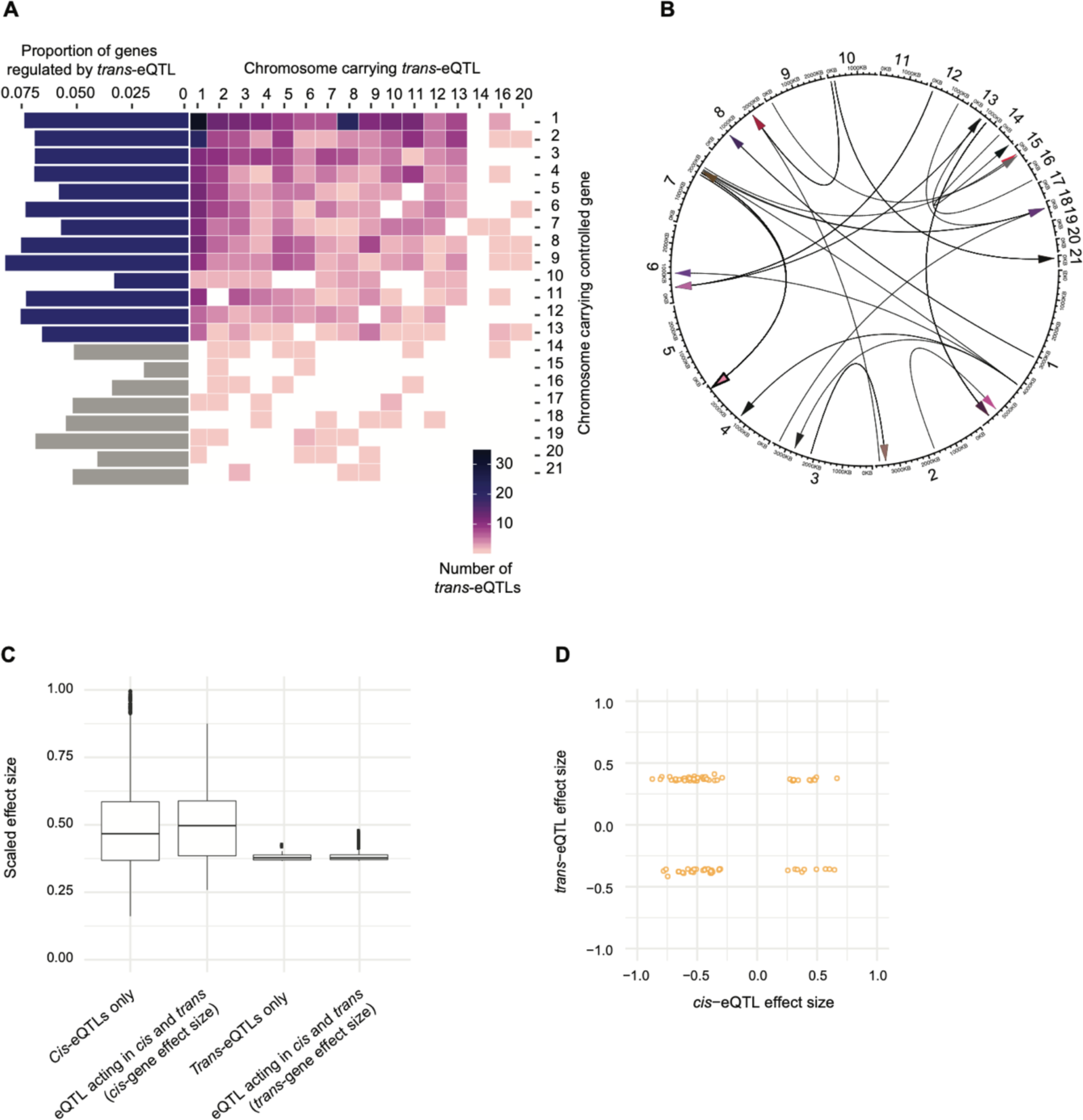
Genome-wide distribution of mapped trans-eQTLs. (A) Distribution of mapped *trans*-eQTLs on core and accessory chromosomes (retained following approximate pass). Barplots on the right indicate the proportion of genes categorized by chromosomes regulated by *trans*-eQTLs. (B) Distribution of *trans*-eQTLs on core and accessory chromosomes. (C) The effect size of overlapping pairs of *cis* and *trans*-eQTLs. (D) Matching of effect sizes for overlapping pairs of cis and *trans*-eQTLs.

### Trait-associated SNPs are overrepresented in cis-regulatory variants

How associated polymorphisms influence phenotypic trait variation remains often unknown. We performed analyses linking known variants associated with virulence and fungicide resistance of the pathogen to mapped *cis*-eQTLs. Trait-associated SNPs were enriched in *cis*-regulatory variants compared to the genomic background with an odds ratio of 20.82 (95% confidence interval 19.16-22.6). Among all analyzed traits, SNPs associated with azole fungicide resistance and reproduction on a wheat cultivar showed the highest degrees of colocalization with mapped *cis*-eQTLs (Figure 6A, Suppl. Figure S6). Furthermore, moderately frequent regulatory variants (minor allele frequency 0.15-0.2 and 0.4-0.45) showed the highest overrepresentation of colocalized trait-associated SNPs (Figure 6B-C). To estimate selection pressure acting on regulatory variants, we assessed potential enrichment of mapped eQTLs for rare or common variants globally. C*is*-eQTLs with low frequency variants were most overrepresented (Figure 7A). We also observed an inverse relationship between the effect size of the *cis*-eQTL and the minor allele frequency of the top associated SNP (Figure 7B). Hence, low frequency variants show the strongest effects on gene expression overall. A skew towards low frequency of high-impact regulatory variants is consistent with purifying selection acting against regulatory variation in the population.

**Figure 6.**
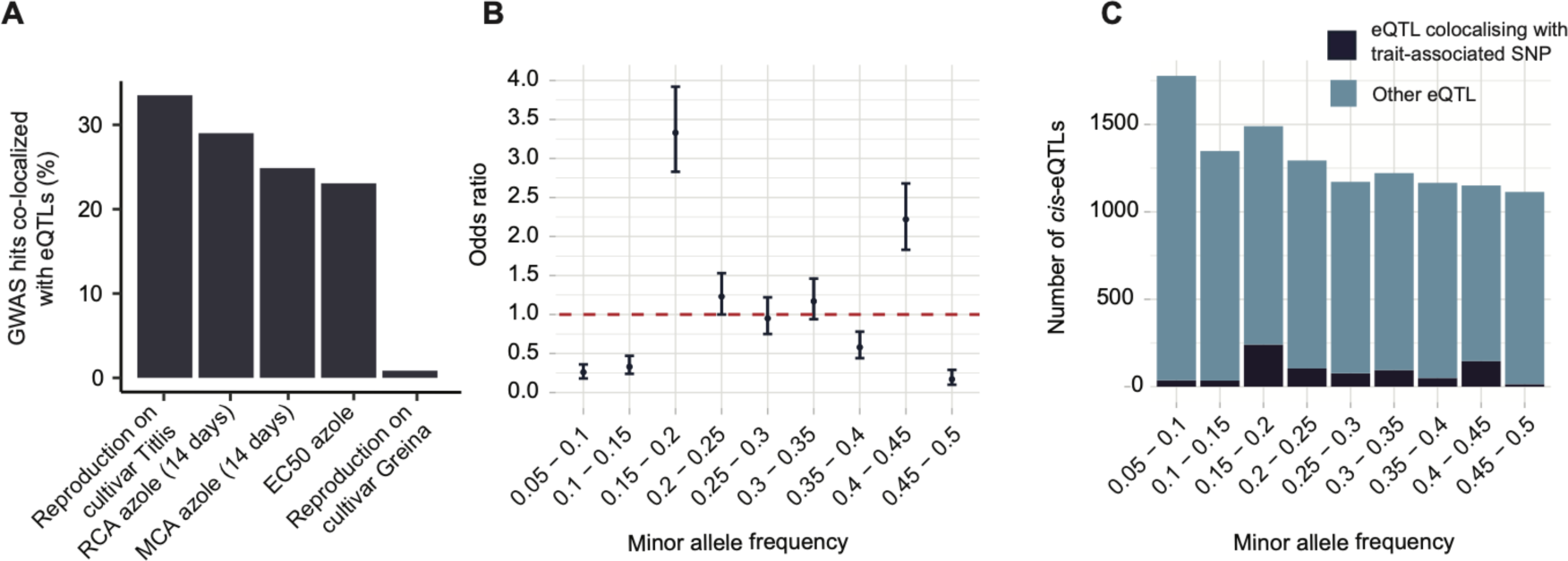
Colocalization of eQTLs with GWAS-associated SNPs. (A) Percentage of phenotypic trait-associated SNPs colocalized with eQTLs. MCA: mean of colony area; RCA: ratio of colony area; EC50: Half-inhibitory concentration. (B) Enrichment of phenotypic trait associated *cis*-eQTLs across the minor allele frequency spectrum. (C) Number of overlapping phenotypic trait-associated SNPs and *cis*-eQTLs in different minor allele frequencies.

**Figure 7.**
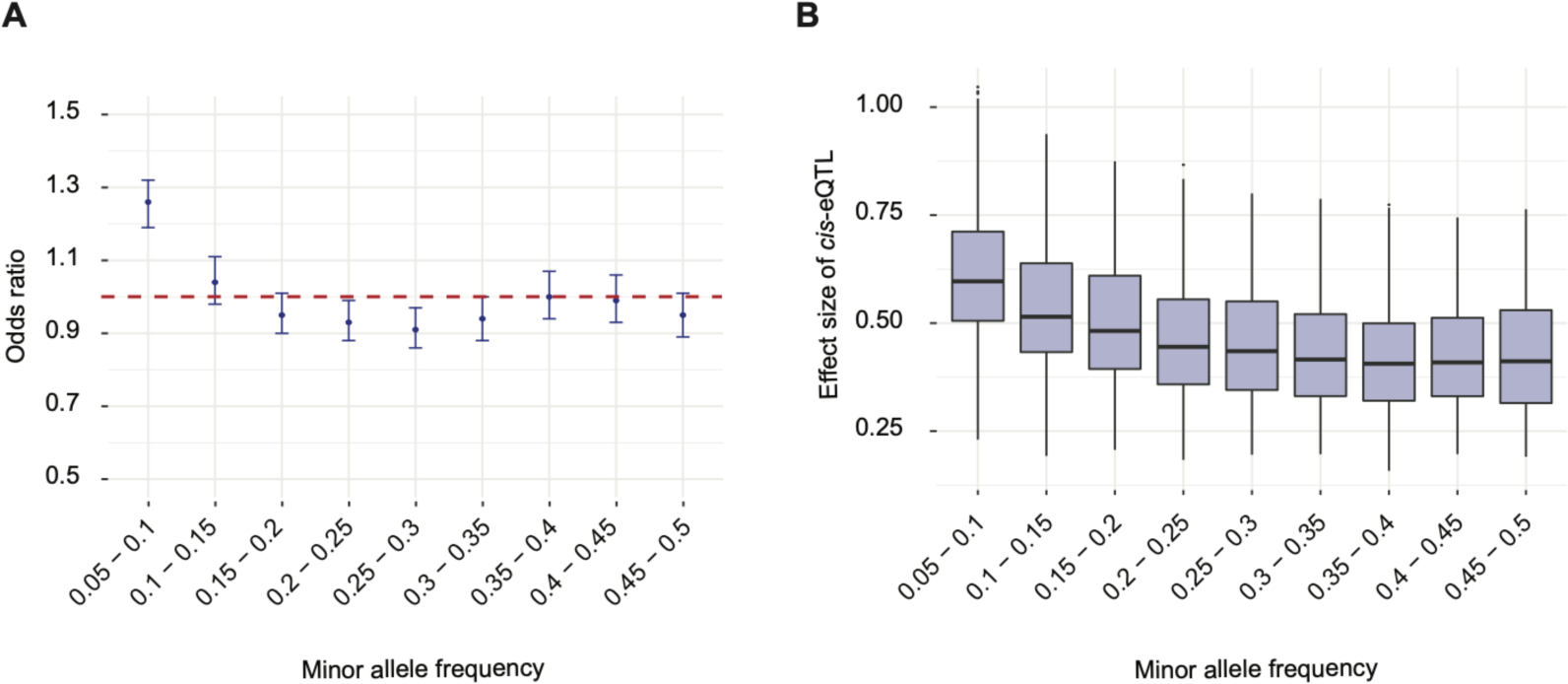
Allele frequency spectrum of *cis*-eQTL variants in the population. (A) Enrichment of *cis*-eQTLs across minor allele frequency categories. (B) Effect size of *cis*-eQTLs across minor allele frequency categories.

### Regulatory architectures of virulence and fungicide sensitivity-associated genes

The eQTL mapping revealed a complex regulatory architecture of previously characterized genes important for the pathogenesis and fungicide resistance (Suppl. Table S4). The expression of small-secreted proteins is coordinated by transcriptional factors and epigenetic regulation. ZtWor1 is an important transcriptional regulator for the expression of small-secreted proteins [46]. ZtWor1 was previously characterized as influencing necrosis and pycnidia development on wheat. We found a *trans*-eQTL in a gene encoding the protein arginine methyltransferase 5 (PRMT5) associated with the expression of the transcriptional regulator ZtWor1. PRMT5 is a methyltransferase for the histone H2A and H4 involved in chromatin remodeling and associated with transcriptional silencing of the gene [47]. Coordinated expression of methyltransferases and the transcriptional regulator of small-secreted proteins may underpin the epigenetic regulation of genes encoding effectors.

Secondary metabolites also play a pivotal role in fungal pathogenesis [48]. The biosynthesis is mediated by coordinated regulation of co-localized genes organized in clusters. A polyketide synthase cluster on chromosome 13 includes a gene encoding the necrosis-inducing protein NPP1 important for virulence in many fungal pathogens even though functional characterization in *Z. tritici* revealed no virulence association [49]. We identified a polymorphism in a gene encoding a fungal trichothecene efflux pump (7_00175) associated with the expression of the gene encoding the necrosis-inducing protein (NPP1; 13_00229). The association supports the idea that the trichothecene efflux pump acts as a transporter in concert with the PKS gene cluster. Azole fungicide sensitivity in *Z. tritici* can be mediated by multiple mechanisms including the efflux of fungicides by specific transporter such as ATP-binding cassette (ABC) transporters [50]. We identified an association of polymorphisms in the gene encoding the ABC transporter MgATR2 with polymorphisms in a gene encoding an MFS, which is a superfamily of membrane transporter proteins. These links underline that ABC transporter efflux functions may contribute to the variation in azole fungicide sensitivity in populations.

## Discussion

We generated the first genome-wide map of regulatory variants underpinning variation in gene expression in a fungal plant pathogen. The majority of all genes in the genome segregated at least one *cis*-eQTL in the mapping population derived from a single field site. Variants upstream of the TSS and untranslated regions were more likely associated with expression variation but with lower effect size. Pathogenicity-related genes were less likely associated to eQTLs consistent with epigenetic regulation playing a prominent role. Phenotypic trait variation is likely governed substantially by gene expression variation.

We successfully identified at least one *cis*-eQTL for two-thirds of all genes indicating how broadly regulatory variation segregates within populations. Genes lacking associated regulatory variants possibly reflect statistical limitations in detecting small effect sizes or the relevant variants were fixed in the analyzed population. *Cis*-eQTLs reported from other intra-species mapping studies varied from 20% of genes in 85 diverse yeast strains [51], 28% of genes in the *D. melanogaster* genetic reference panel (DGRP) of 39 inbred lines [12], 13% of genes in *A. thaliana* populations [52], 6.7% of genes in humans with 270 individuals from multiple populations and 31% of the genes in 801 European and 1,032 African Americans [53–55]. Mapping studies differed by panel size and genetic diversity of the mapping population. The comparatively smaller population size used for mapping might explain the lower proportion of *cis*-eQTLs in the populations of *A. thaliana* and DGRP lines of *D. melanogaster* (a representative sample of naturally segregating genetic variation). The small number of eQTLs mapped in the comparatively large mapping population of *S. cerevisiae* may stem from slow linkage disequilibrium decay. Population genomics studies in *Z. tritici* reported that ∼90% of the global genetic variation is found within single populations and with few indications of bottlenecks [36, 37, 56–58]. Hence, *Z. tritici* populations are highly amenable to the efficient mapping of *cis*-eQTLs.

A substantial minority of all genes were mapped with multiple eQTLs consistent with the high-resolution of the panel and complexity of *cis*-regulatory variation in the species. The primary regulatory variant for each gene was located close to the TSS and the effect of regulatory variants on gene expression was inversely proportional to the distance of the variant from the TSS. Hence, dominant regulatory variants are likely binding sites and promoters proximal to TSS. Indels in introns, exons, and 3’ UTR had stronger effects on gene expression compared to SNPs. Indels are more likely to disrupt the integrity of splice sites, RNA binding protein motifs in 3’UTR and are, hence, more likely causal variants for gene expression variation than SNPs [59]. Indel mutations at DNA binding sites can strongly reduce the binding affinity of DNA binding proteins such as TFs, whereas the degenerate nature of DNA binding sites can balance out more easily the effects of point mutations [60]. Intronic polymorphisms had stronger effects on gene expression variation but were less likely to be identified as *cis*-eQTLs, which is consistent with stronger selective constraints on indel polymorphisms. In plants such as *A. thaliana*, the larger introns can likely more effectively buffer deleterious effects of indel mutations [61]. Also, polymorphism in introns may result in alternatively spliced mRNA and functionally diverse proteins with possible phenotypic consequences [62]. The generalist plant pathogen *Sclerotinia sclerotiorum* was found to accumulate alternative transcript isoforms upon infection depending on the host identity, which could indicate alternative splicing events are taking place upon infection to generate functionally diverse secreted proteins [63]. Hence, the strength of constraints on regulatory or splice variants in coding sequences has likely consequences for the evolvability of gene regulation within the species.

Gene regulatory polymorphisms were unevenly distributed among functional categories of genes. eQTLs were underrepresented in genes encoding functions important for pathogenesis *i.e.*, effectors and secondary metabolites. This observation may stem from unequal success in variant calling steps since genes encoding effector proteins tend to localize in the vicinity of TEs [64]. The repetitive nature of TEs may prevent the identification of reliable SNPs, which in turn could lead to the underestimation of *cis*-eQTLs. In contrast to the lower proportion of genes mapped with eQTLs, the likelihood of a variant being called an eQTL is higher for variants close to the genes encoding effectors, secondary metabolites, and genes highly expressed during plant infection. However, effect sizes of these *cis-* eQTLs were comparatively low to eQTLs for other gene categories. This could suggest that gene regulation for pathogenicity-associated genes tends to be mediated by epigenetics rather than *cis*-regulatory elements. Epigenetic gene regulation can form long-range chromatin interactions mediated by chromatin state changes and can regulate the expression of the target gene from hundreds of kilobases away. These chromatin interactions are physical interactions of DNA mediated by histone methylation marks and do not require nucleotide variants [65]. This observation suggests the involvement of epigenetic regulation as a major player in effector gene regulation consistent with studies showing effector expression being influenced by TEs [66, 67].

Our *trans-*eQTL mapping identified only few genes associated with regulatory variation compared to *cis*-eQTLs. A similar observation was reported in humans (0.3 % of the genes associated with *trans-* eQTLs) [68] and recombinant inbred lines of *D. melanogaster* [69], whereas a study of 1,012 yeast segregants from a cross between a laboratory and a wine strain reported almost all the expressed genes as having at least one *trans-*eQTL (Albert et al., 2018a). A lower number of genes mapped with *trans-* eQTLs than *cis*-eQTLs might be due to the lower mapping power resulting from the high multiple testing burden in *trans-*eQTL analyses. Unlike the mapping studies in *S. cerevisiae* [21] and maize [71], we identified no *trans-*regulatory hot spots (*i.e. trans-*eQTL regulating large gene sets). We expected core and accessory chromosomes of *Z. tritici* to show regulatory links, however we found that core chromosome genes were almost exclusively regulated core chromsome *trans-*eQTLs. This compartmentalization is consistent with distinct organizations of core and accessory chromosomes. Studies on the filamentous fungus *Epichloë festucae* [72] and *D. melanogaster* [73] both underlined the that the 3D structure of the genome can mediate transcription and regulate gene expression. Hence, the *Z. tritici* 3D chromosomal conformation may allow for only few core-accessory chromosome links.

The *trans-*eQTLs mapped for virulence and fungicide sensitivity-associated genes suggested the existence of complex regulatory networks. The *trans-*regulatory polymorphism in the gene encoding a methyltransferase (PRMT5) is associated with expression variation of the transcription factor known as a positive regulator of major virulence genes in the pathogen. PRMT5 mediates the methylation of histone arginine and plays an important role in chromatin dynamics [47]. The coordinated regulation of histone methyltransferase and the regulator of virulence genes highlights the importance of the epigenetic regulation layer of effector gene regulation. The genes encoding secondary metabolites tend to colocalize in clusters containing one or more core biosynthetic genes, accessory genes, major regulators, and transporters [74, 75]. In our *trans-*eQTL mapping, we identified a polymorphism in an exon of the gene encoding a trichothecene efflux pump and regulating the expression of the gene encoding the necrosis-inducing protein (NPP1). NPP1 is also a predicted effector suggesting that secondary metabolite gene clusters may be capable of coordinating gene regulation outside of the cluster. Such effects may include the regulation of transporters relevant for the transport of the secondary metabolite.

Gaining a mechanistic understanding how polymorphism identified from phenotype-genotype association mapping contributes to phenotypic variation is often challenging. Colocalization of eQTLs and trait-associated SNPs enables more focused hypothesis-testing. Our colocalization approach using SNPs associated with virulence and fungicide sensitivity identified that phenotypic trait-associated SNPs were enriched for regulatory polymorphisms with a quarter of the SNPs colocalizing with mapped *cis*-eQTLs. This underlines that regulatory polymorphism is likely playing a major role in phenotypic trait variation. Hence, recent adaptation of the pathogen to hosts and the environment may have been achieved with substantial contributions from regulatory polymorphisms. Ascertaining standing variation for genetic and regulatory variants enables more precise predictions of the evolutionary potential of species.

## Materials and methods

### Library preparation, genome and transcriptome sequencing

Isolates of *Z. tritici* were collected from an experimental wheat field planted with different cultivars [36, 76] and grown for 10 days in yeast-sucrose broth (YSB) at 18°C. Total genomic DNA was extracted using the QIAGEN DNAeasy Plant Mini Kit and the Illumina library was prepared using a TruSeq Nano DNA Library kit (Illumina, Inc.). Libraries with an insert size of ∼550 bp were sequenced for a read length of 100 bp in paired-end mode on a HiSeq 4000 at the iGE3 sequencing platform (Geneva, Switzerland). For RNA sequencing, the same isolates were cultured in a Vogel’s Medium N (Minimal) [77] modified as ammonium nitrate replaced with potassium nitrate and ammonium phosphate [78] without sucrose and agarose to induce hyphal growth. Total RNA was isolated from the filtered mycelium after 10 -15 days using the NucleoSpin® RNA Plant and Fungi kit. RNA concentration and integrity were checked using a Qubit 2.0 Fluorometer and an Agilent 4200 TapeStation System, respectively. Only high-quality RNA (RIN>8) was used to prepare TruSeq stranded mRNA libraries with a 150 bp insert size and sequenced on an Illumina HiSeq 4000 in the single-end mode for 100 bp.

### Variant calling and filtering

DNA sequences were checked for quality using FastQC version 0.11.5 [79] and trimmed with trimmomatic version 0.36 [80] to remove adapter sequences and low-quality reads with parameters ILLUMINACLIP: TruSeq3-PE.fa:2:30:10 LEADING:3 TRAILING:3 SLIDING WINDOW:4:15 MINLEN:36. Trimmed sequences were aligned to the *Z. tritici* reference genome of IPO323 [81] and mitochondrial sequence (accession EU090238.1) using Bowtie2 version 2.3.4.3 [82] with the option -- very-sensitive-local. Aligned sequences were used for variant calling with the HaplotypeCaller integrated in the Genome Analysis Toolkit (GATK) v. 4.0.11.0. [83]. SNPs and indels were separated using the SelectVariants tool. SNPs and indels were quality filtered with the VariantFiltration tool. We retained SNPs with QUAL>1000, AN=20, QD>5.0, MQ>20.0, as well as ReadPosRankSum, MQRankSum, BaseQRankSum between -2.0 and 2.0. Variants passing the quality filtration were further filtered to remove multiallelic sites using the bcftools (version 1.9) --norm option. After that, the variants were filtered to keep only sites genotyped in at least 90% of the individuals and rare variants (< 5%) were removed using VCFtools [84] and the --max-missing option and bcftools (version 1.9) [85] -q 0.05: minor option.

### RNA-seq analyses

RNA-seq data were checked for quality using FastQC (version 0.11.5) [79] and trimmed with trimmomatic version 0.36 [80] to remove adapter sequences and low-quality reads with parameters: ILLUMINACLIP: TruSeq3-SE.fa:2:30:10 LEADING:3 TRAILING:3 SLIDING WINDOW:4:15 MINLEN:36. Trimmed sequences were aligned to the *Z. tritici* reference genome of IPO323 [81] using HISAT2 [86] (version 2.1.0) with the parameter “--RNA-strandedness reverse”.

*De novo* trans*criptome assembly and UTR prediction*

For *de novo* transcriptome assembly, we used RNA-seq datasets generated for the reference isolate IPO323 from different plant infection stages (1, 4, 9, 14 and 21 days post infection) and culture media (Czapek-Dox broth, potato dextrose broth) available on NCBI (accession numbers: ERS684130-37, ERS684123-29, ERS683735-40, ERS6837343). We used also Illumina RNA-seq reads generated from minimal media grown following the above-described methods. For *in planta* RNA-seq data, reads were filtered by aligning to the IPO323 reference genome [81] to remove transcripts from wheat. Trinity (version v2.8.3) [87] was used for *de novo* transcriptome assembly using the --jaccard_clip parameter to reduce the occurrence of fused transcripts in dense fungal genomes. We performed *de novo* assembly using different sets of RNA-seq samples including up to a maximum of ∼5 million reads per dataset. Next, we used GenomeThreader version 1.7.1 [88] to align *de novo* assembled transcripts to the reference genome requiring a minimum alignment score of 0.98. Along with *de novo* assembled transcripts, we used Iso-Seq reads generated for IPO323 to improve UTR prediction [89]. We then checked for overlaps of d*e novo* assembled transcripts and Iso-Seq reads with IPO323 gene models [90] using bedtools (version v2.27.1)[91]. Overlapping transcript sequences were further filtered to remove transcripts overlapping more than one gene and transcripts smaller than the predicted gene model. After filtering, the start and end position of the longest predicted transcript was considered as the 5’UTR start and 3’UTR end of each gene.

### Mapping cis-eQTL using permutations

We used QTLtools (version 1.1) [92] for transcriptomic data filtering and the mapping of eQTL. Reads mapped to gene models [90] were counted with the QTLtools --quan mode. Only reads with a minimum Phred mapping quality >10 was kept for further analyses. Normalization of the counts was done using the --rpkm option implemented in the QTLtools --quan mode. To determine the optimal configuration for eQTL discovery, we performed eQTL mapping with 1000 permutations and 5kb *cis* windows filtering for genes with RPKM=0. Principal components (PCs) explaining variance at the genotype and expression level were calculated using the --PCA mode in QTLtools. To determine the number of PCs for population structure correction and technical variance in the dataset, independent eQTL analyses were performed in the *cis* --permutation mode with 1000 permutations and 5kb *cis* windows with differing numbers of PCs. The permutation *p-*values are false discovery rate (FDR) corrected to identify the top - eQTL significant at a 5% FDR level. To map *cis*-eQTLs with an independent effect on gene expression, we used the QTLtools --*cis* conditional option and reported 5% FDR corrected eQTLs in association with the top variant reported from --*cis* permutation analyses. For this, we have chosen a *cis* window of 10kb equidistant from the TSS.

### Mapping trans-eQTL using permutations

To map trans eQTLs we used the --trans mode in QTLtools (version 1.1). We ran the full pass mode to identify the top candidates of trans eQTLs. The full permutation scheme in the QTLtools trans analysis permutes all expression phenotypes and genetic variants excluding the window defined for *cis* variants. We ran the nominal pass with the threshold of 1e-5 excluding variants in the window of 200 kb equidistant from the start codon of each focal gene. Multiple testing correction was performed using a full permutation scheme with 100 permutations. We have retained trans eQTLs if the association was significant at a 5% FDR in at least 80% of the permutations. The approximation pass selected 1,000 expression phenotypes randomly, permutated expression phenotypes and tested for associations against all variants as a less stringent threshold to retain *trans* eQTL.

### Colocalizing mapped eQTLs with GWAS-associated variants

To assess colocalization of eQTLs with GWAS signals, we used the RTC mode in QTLtools (version 1.1) [92]. We chose variants associated with phenotypes from a GWAS study based on a mapping population established from the same wheat field [93]. We analyzed GWAS for fungicide resistance, virulence, and reproduction of the pathogen for co-localization with mapped *cis*-eQTLs. We retained co-localized variants if the RTC score was >0.9 and the linkage disequilibrium (*r*^2^) between the variants >0.5.

## Acknowledgments

We are grateful for helpful comments by Thomas Badet and Sabina Tralamazza on a previous version of this manuscript. Nicolas Lapalu and Marc-Henri Lebrun facilitated access to sequencing read datasets. Sequencing data was generated using facilities of the Genetic Diversity Centre (GDC), ETH Zurich, and the Institute of Genetics and Genomics, Geneva.

## References

1. Ladunga I. Computational Biology of Transcription Factor Binding. Human Press; 2010.

2. Harbison CT, Gordon DB, Lee TI, Rinaldi NJ, Macisaac KD, Danford TW, et al. Transcriptional regulatory code of a eukaryotic genome. Nature. 2004;431:99–104.

3. Tjian R. The biochemistry of transcription in eukaryotes: A paradigm for multisubunit regulatory complexes. Philosophical Transactions of the Royal Society B: Biological Sciences. 1996;351:491–9.

4. Birney E, Stamatoyannopoulos JA, Dutta A, Guigó R, Gingeras TR, Margulies EH, et al. Identification and analysis of functional elements in 1% of the human genome by the ENCODE pilot project. Nature. 2007;447:799–816.

5. Willbanks A, Leary M, Greenshields M, Tyminski C, Heerboth S, Lapinska K, et al. The Evolution of Epigenetics: From Prokaryotes to Humans and Its Biological Consequences. Genet Epigenet. 2016;8:25.

6. Kita R, Venkataram S, Zhou Y, Fraser HB, Rine J. High-resolution mapping of cis-regulatory variation in budding yeast. Proc Natl Acad Sci U S A. 2017. https://doi.org/10.1073/pnas.1717421114.

7. Salinas F, De Boer CG, Abarca V, García V, Cuevas M, Araos S, et al. Natural variation in non-coding regions underlying phenotypic diversity in budding yeast. Sci Rep. 2016;6:1–13.

8. Bailey SF, Hinz A, Kassen R. Adaptive synonymous mutations in an experimentally evolved *Pseudomonas fluorescens* population. Nat Commun. 2014;5:1–7.

9. Kita R, Venkataram S, Zhou Y, Fraser HB. High-resolution mapping of cis-regulatory variation in budding yeast. Proc Natl Acad Sci U S A. 2017;114:E10736–44.

10. Thompson DA, Cubillos FA. Natural gene expression variation studies in yeast. Yeast. 2017;34:3– 17.

11. Lutz S, Brion C, Kliebhan M, Albert FW. DNA variants affecting the expression of numerous genes in trans have diverse mechanisms of action and evolutionary histories. PLoS Genet. 2019;15.

12. Massouras A, Waszak SM, Albarca-Aguilera M, Hens K, Holcombe W, Ayroles JF, et al. Genomic Variation and Its Impact on Gene Expression in *Drosophila melanogaster*. PLoS Genet. 2012;8:e1003055–14.

13. Osada N, Miyagi R, Takahashi A. Cis- and Trans-regulatory Effects on Gene Expression in a Natural Population of *Drosophila melanogaster*. Genetics. 2017;206:2139–48.

14. Enard D, Messer PW, Petrov DA. Genome-wide signals of positive selection in human evolution. Genome Res. 2014;24:885–95.

15. Schaub MA, Boyle AP, Kundaje A, Batzoglou S, Snyder M. Linking disease associations with regulatory information in the human genome. Genome Res. 2012;22:1748–59.

16. Vandiedonck C. Genetic association of molecular traits: A help to identify causative variants in complex diseases. Clin Genet. 2018;93:520–32.

17. Snoek BL, Sterken MG, Bevers RPJ, Volkers RJM, Hof A van’t, Brenchley R, et al. Contribution of trans regulatory eQTL to cryptic genetic variation in C. elegans. BMC Genomics. 2017;18:1–15.

18. Rigau M, Juan D, Valencia A, Rico D. Intronic CNVs and gene expression variation in human populations. PLoS Genet. 2019;15:e1007902.

19. Uzunović J, Josephs EB, Stinchcombe JR, Wright SI, Parsch J. Transposable Elements Are Important Contributors to Standing Variation in Gene Expression in *Capsella Grandiflora*. Mol Biol Evol. 2019;36:1734–45.

20. Josephs EB, Lee YW, Stinchcombe JR, Wright SI. Association mapping reveals the role of purifying selection in the maintenance of genomic variation in gene expression. Proc Natl Acad Sci U S A. 2015;112:15390.

21. Albert FW, Bloom JS, Siegel J, Day L, Kruglyak L. Genetics of trans-regulatory variation in gene expression. Elife. 2018;7:204–20.

22. Soltis NE, Caseys C, Zhang W, Corwin JA, Atwell S, Kliebenstein DJ. Pathogen genetic control of transcriptome variation in the arabidopsis thaliana - *Botrytis cinerea* pathosystem. Genetics. 2020;215:253–66.

23. Chang J, Au CH, Cheng CK, Kwan HS. eQTL network analysis reveals that regulatory genes are evolutionarily older and bearing more types of PTM sites in Coprinopsis cinerea. 2018;:1–21.

24. Wu VW, Thieme N, Huberman LB, Dietschmann A, Kowbel DJ, Lee J, et al. The regulatory and transcriptional landscape associated with carbon utilization in a filamentous fungus. Proc Natl Acad Sci U S A. 2020;117:6003–13.

25. Omrane S, Audéon C, Ignace A, Duplaix C, Aouini L, Kema G, et al. Plasticity of the MFS1 Promoter Leads to Multidrug Resistance in the Wheat Pathogen Zymoseptoria tritici. mSphere. 2017;2.

26. Kretschmer M, Leroch M, Mosbach A, Walker A-S, Fillinger S, Mernke D, et al. Fungicide-Driven Evolution and Molecular Basis of Multidrug Resistance in Field Populations of the Grey Mould Fungus Botrytis cinerea. PLoS Pathog. 2009;5:e1000696.

27. Zhang Q, Ma C, Zhang Y, Gu Z, Li W, Duan X, et al. A single-nucleotide polymorphism in the promoter of a hairpin rna contributes to alternaria alternata leaf spot resistance in apple (Malus × domestica). Plant Cell. 2018;30:1924–42.

28. Palma-Guerrero J, Torriani SFF, Zala M, Carter D, Courbot M, Rudd JJ, et al. Comparative transcriptomic analyses of Zymoseptoria tritici strains show complex lifestyle transitions and intraspecific variability in transcription profiles. Mol Plant Pathol. 2016. https://doi.org/10.1111/mpp.12333.

29. Palma-Guerrero J, Ma X, Torriani SFF, Zala M, Francisco CS, Hartmann FE, et al. Comparative Transcriptome Analyses in Zymoseptoria tritici Reveal Significant Differences in Gene Expression Among Strains During Plant Infection. MPMI. 2017;30:231–44.

30. Soyer JL, El Ghalid M, Glaser N, Ollivier B, Linglin J, Grandaubert J, et al. Epigenetic control of effector gene expression in the plant pathogenic fungus Leptosphaeria maculans. PLoS Genet. 2014;10:e1004227.

31. Tang B, Yan X, Ryder LS, Bautista MJA, Cruz-Mireles N, Soanes DM, et al. Rgs1 is a regulator of effector gene expression during plant infection by the rice blast fungus Magnaporthe oryzae. Proceedings of the National Academy of Sciences. 2023;120:e2301358120.

32. Sánchez-Vallet A, Fouché S, Fudal I, Hartmann FE, Soyer JL, Tellier A, et al. The Genome Biology of Effector Gene Evolution in Filamentous Plant Pathogens. Annu Rev Phytopathol. 2018. https://doi.org/10.1146/annurev-phyto-080516.

33. de Jonge R, Thomma BPHJ. Fungal LysM effectors: extinguishers of host immunity? Trends Microbiol. 2009;17:151–7.

34. Lanver D, Müller AN, Happel P, Schweizer G, Haas FB, Franitza M, et al. The Biotrophic Development of Ustilago maydis Studied by RNA-Seq Analysis. Plant Cell. 2018;30:300–23.

35. Mentlak TA, Kombrink A, Shinya T, Ryder LS, Otomo I, Saitoh H, et al. Effector-Mediated Suppression of Chitin-Triggered Immunity by Magnaporthe oryzae Is Necessary for Rice Blast Disease. Plant Cell. 2012;24:322–35.

36. Singh NK, Karisto P, Croll D. Population-level deep sequencing reveals the interplay of clonal and sexual reproduction in the fungal wheat pathogen Zymoseptoria tritici. Microb Genom. 2021;7:678.

37. Feurtey A, Lorrain C, McDonald MC, Milgate A, Solomon PS, Warren R, et al. A thousand-genome panel retraces the global spread and adaptation of a major fungal crop pathogen. Nature Communications 2023 14:1. 2023;14:1–15.

38. Mcdonald BA, Linde C. Pathogen Population Genetics, Evolutionary Potential, And Durable Resistance. Annu Rev Phytopathol. 2002;40:349–79.

39. Zhan J, Pettway RE, McDonald BA. The global genetic structure of the wheat pathogen Mycosphaerella graminicola is characterized by high nuclear diversity, low mitochondrial diversity, regular recombination, and gene flow. Fungal Genetics and Biology. 2003;38:286–97.

40. Testa A, Oliver R, Hane J. Overview of genomic and bioinformatic resources for Zymoseptoria tritici. Fungal Genetics and Biology. 2015;79:13–6.

41. Fouché S, Badet T, Oggenfuss U, Plissonneau C, Francisco CS, Croll D. Stress-Driven Transposable Element De-repression Dynamics and Virulence Evolution in a Fungal Pathogen. Mol Biol Evol. 2020;37:221–39.

42. Meile L, Croll D, Brunner PC, Plissonneau C, Hartmann FE, McDonald BA, et al. A fungal avirulence factor encoded in a highly plastic genomic region triggers partial resistance to septoria tritici blotch. New Phytologist. 2018;219:1048–61.

43. Krishnan P, Meile L, Plissonneau C, Ma X, Hartmann FE, Croll D, et al. Transposable element insertions shape gene regulation and melanin production in a fungal pathogen of wheat. BMC Biol. 2018;16:78.

44. Abraham LN, Oggenfuss U, Croll D. Population-level transposable element expression dynamics influence trait evolution in a fungal crop pathogen. bioRxiv. 2023;:2023.03.29.534750.

45. Oggenfuss U, Croll D. Recent transposable element bursts are associated with the proximity to genes in a fungal plant pathogen. PLoS Pathog. 2023;19:e1011130.

46. Mirzadi Gohari A, Mehrabi R, Robert O, Ince IA, Boeren S, Schuster M, et al. Molecular characterization and functional analyses of ZtWor1, a transcriptional regulator of the fungal wheat pathogen Zymoseptoria tritici. Mol Plant Pathol. 2013;15:394–405.

47. Litt M, Qiu Y, Huang S. Histone arginine methylations: Their roles in chromatin dynamics and transcriptional regulation. Bioscience Reports. 2009;29:131–41.

48. Macheleidt J, Mattern DJ, Fischer J, Netzker T, Weber J, Schroeckh V, et al. Regulation and Role of Fungal Secondary Metabolites. Annual Review of Genetics. 2016;50:371–92.

49. Motteram J, Küfner I, Deller S, Brunner F, Hammond-Kosack KE, Nürnberger T, et al. Molecular characterization and functional analysis of MgNLP, the sole NPP1 domain-containing protein, from the fungal wheat leaf pathogen mycosphaerella graminicola. Molecular Plant-Microbe Interactions. 2009;22:790–9.

50. Zwiers LH, Stergiopoulos I, Van Nistelrooy JGM, De Waard MA. ABC transporters and azole susceptibility in laboratory strains of the wheat pathogen Mycosphaerella graminicola. Antimicrob Agents Chemother. 2002;46:3900–6.

51. Kita R, Venkataram S, Zhou Y, Fraser H. High-Resolution Mapping Of cis-Regulatory Variation In Budding Yeast. 2018;:1–62.

52. Zan Y, Shen X, Forsberg SKG, Carlborg Ö. Genetic Regulation of Transcriptional Variation in Natural *Arabidopsis thaliana* Accessions. G3: Genes|Genomes|Genetics. 2016;6:2319–28.

53. Shang L, Smith JA, Zhao W, Kho M, Turner ST, Mosley TH, et al. Genetic Architecture of Gene Expression in European and African Americans: An eQTL Mapping Study in GENOA. Am J Hum Genet. 2020;106:496–512.

54. Stranger BE, Montgomery SB, Dimas AS, Parts L, Stegle O. Patterns of Cis Regulatory Variation in Diverse Human Populations. PLoS Genet. 2012;8:1002639.

55. Stranger BE, Nica AC, Forrest MS, Dimas A, Bird CP, Beazley C, et al. Population genomics of human gene expression. Nat Genet. 2007;39:1217–24.

56. Morais D, Duplaix C, Sache I, Laval V, Suffert F, Walker A-S. Overall stability in the genetic structure of a Zymoseptoria tritici population from epidemic to interepidemic stages at a small spatial scale. Eur J Plant Pathol. 2019;154:423–36.

57. Schnieder F, Koch G, Jung C, Verreet J-A. Genotypic diversity of the wheat leaf blotch pathogen Mycosphaerella graminicola (anamorph) Septoria tritici in Germany. 2001.

58. McDonald BA, Suffert F, Bernasconi A, Mikaberidze A. How large and diverse are field populations of fungal plant pathogens? The case of Zymoseptoria tritici. Evol Appl. 2022;15:1360–73.

59. Huang J, Chen J, Esparza J, Ding J, Elder JT, Abecasis GR, et al. eQTL mapping identifies insertion- and deletion-specific eQTLs in multiple tissues. Nat Commun. 2015;6:6821.

60. Deplancke B, Alpern D, Gardeux V. The Genetics of Transcription Factor DNA Binding Variation. Cell. 2016;166:538–54.

61. Mukherjee D, Saha D, Acharya D, Mukherjee A, Chakraborty S, Ghosh TC. The role of introns in the conservation of the metabolic genes of Arabidopsis thaliana. Genomics. 2018;110:310–7.

62. Lalonde E, Ha KCH, Wang Z, Bemmo A, Kleinman CL, Kwan T, et al. RNA sequencing reveals the role of splicing polymorphisms in regulating human gene expression. Genome Res. 2011;21:545– 54.

63. Ibrahim HMM, Kusch S, Didelon M, Raffaele S. Genome-wide alternative splicing profiling in the fungal plant pathogen Sclerotinia sclerotiorum during the colonization of diverse host families. bioRxiv. 2020;:2020.05.13.094565.

64. Torres DE, Oggenfuss U, Croll D, Seidl MF. Genome evolution in fungal plant pathogens: looking beyond the two-speed genome model. Fungal Biology Reviews. 2020;34:136–43.

65. Yu J, Hu M, Li C. Joint analyses of multi-tissue Hi-C and eQTL data demonstrate close spatial proximity between eQTLs and their target genes. BMC Genet. 2019;20:43.

66. Fouché S, Badet T, Oggenfuss U, Plissonneau C, Francisco CS, Croll D. Stress-Driven Transposable Element De-repression Dynamics and Virulence Evolution in a Fungal Pathogen. Mol Biol Evol. 2020;37.

67. Abraham LN, Oggenfuss U, Croll D. Population-level transposable element expression dynamics influence trait evolution in a fungal crop pathogen. bioRxiv. 2023;:2023.03.29.534750.

68. Bryois J, Buil A, Evans DM, Kemp JP, Montgomery SB. Cis and Trans Effects of Human Genomic Variants on Gene Expression. PLOS Genet. 2014;10:1004461.

69. Genissel A, McIntyre LM, Wayne ML, Nuzhdin S V. Cis and Trans Regulatory Effects Contribute to Natural Variation in Transcriptome of Drosophila melanogaster. Mol Biol Evol. 2008;25:101–10.

70. Albert FW, Bloom JS, Siegel J, Day L, Kruglyak L. Genetics of trans-regulatory variation in gene expression. Elife. 2018;7:1–39.

71. Wang X, Chen Q, Wu Y, Lemmon ZH, Xu G, Huang C, et al. Genome-wide Analysis of Transcriptional Variability in a Large Maize-Teosinte Population. Mol Plant. 2018;11:443–59.

72. Id DJW, Ganley ARD, Id CAY, Liachko Id I, Schardlid CL, Dupont P-Y, et al. Repeat elements organise 3D genome structure and mediate transcription in the filamentous fungus Epichloë festucae. 2018. https://doi.org/10.1371/journal.pgen.1007467.

73. Cubenãs-Potts C, Rowley MJ, Lyu X, Li G, Lei EP, Corces VG. Different enhancer classes in Drosophilabind distinct architectural proteins and mediate unique chromatin interactions and 3D architecture. Nucleic Acids Res. 2017;45:1714–30.

74. Cairns T, Meyer V. In silico prediction and characterization of secondary metabolite biosynthetic gene clusters in the wheat pathogen Zymoseptoria tritici. BMC Genomics. 2017;18:1–16.

75. Hassani MA, Oppong-Danquah E, Feurtey A, Tasdemir D, Stukenbrock EH. Differential Regulation and Production of Secondary Metabolites among Isolates of the Fungal Wheat Pathogen Zymoseptoria tritici. Appl Environ Microbiol. 2022;88.

76. Karisto P, Dora S, Mikaberidze A. Measurement of infection efficiency of a major wheat pathogen using time-resolved imaging of disease progress. Plant Pathol. 2019;68:163–72.

77. Vogel HJ, Vogel J, Vogel H, Vogel H. A convenient growth medium for Neurospora crassa. 1956.

78. Perkins DD. How to choose and prepare media. 2006;:1–15.

79. S Andrews. FastQC: a quaFastQC: a quality control tool for high throughput sequence datality control tool for high throughput sequence data. Babraham Bioinformatics, Babraham. 2010.

80. Bolger AM, Lohse M, Usadel B. Trimmomatic: A flexible trimmer for Illumina sequence data. Bioinformatics. 2014;30:2114–20.

81. Goodwin SB, M’Barek S ben, Dhillon B, Wittenberg AHJ, Crane CF, Hane JK, et al. Finished genome of the fungal wheat pathogen Mycosphaerella graminicola reveals dispensome structure, chromosome plasticity, and stealth pathogenesis. PLoS Genet. 2011;7.

82. Langmead B, Salzberg SL. Fast gapped-read alignment with Bowtie 2. Nat Methods. 2012;9:357.

83. van der Auwera G, O’Connor B, Safari an OMCompany. Genomics in the Cloud: Using Docker, GATK, and WDL in Terra. Genomics in the Cloud. 2020;:300.

84. Danecek P, Auton A, Abecasis G, Albers CA, Banks E, DePristo MA, et al. The variant call format and VCFtools. Bioinformatics. 2011;27:2156–8.

85. Danecek P, Bonfield JK, Liddle J, Marshall J, Ohan V, Pollard MO, et al. Twelve years of SAMtools and BCFtools. Gigascience. 2021;10.

86. Zhang Y, Park C, Bennett C, Thornton M, Kim D. Rapid and accurate alignment of nucleotide conversion sequencing reads with HISAT-3N. Genome Res. 2021;31:1290–5.

87. Grabherr MG, Haas BJ, Yassour M, Levin JZ, Thompson DA, Amit I, et al. Trinity: reconstructing a full-length transcriptome without a genome from RNA-Seq data. Nat Biotechnol. 2011;29:644.

88. Gremme G, Brendel V, Sparks ME, Kurtz S. Engineering a software tool for gene structure prediction in higher organisms. Inf Softw Technol. 2005;47:965–78.

89. Lapalu N, Lamothe L, Petit Y, Genissel A, Delude C, Feurtey A, et al. Improved gene annotation of the fungal wheat pathogen Zymoseptoria tritici based on combined Iso-Seq and RNA-Seq evidence. bioRxiv. 2023;:2023.04.26.537486.

90. Grandaubert J, Bhattacharyya A, Stukenbrock EH. RNA-seq-Based Gene Annotation and Comparative Genomics of Four Fungal Grass Pathogens in the Genus Zymoseptoria Identify Novel Orphan Genes and Species-Specific Invasions of Transposable Elements. G3 (Bethesda). 2015;5:1323–33.

91. Quinlan AR, Hall IM. BEDTools: a flexible suite of utilities for comparing genomic features. Bioinformatics. 2010;26:841–2.

92. Delaneau O, Ongen H, Brown AA, Fort A, Panousis NI, Dermitzakis ET. A complete tool set for molecular QTL discovery and analysis. Nat Commun. 2017;8:15452–7.

93. Dutta A, Hartmann FE, Francisco CS, McDonald BA, Croll D. Mapping the adaptive landscape of a major agricultural pathogen reveals evolutionary constraints across heterogeneous environments. bioRxiv. 2020;:2020.07.30.229708.

