## Supplementary Figures for "Genome-wide expression QTL mapping reveals the highly dynamic regulatory landscape of a major wheat pathogen"

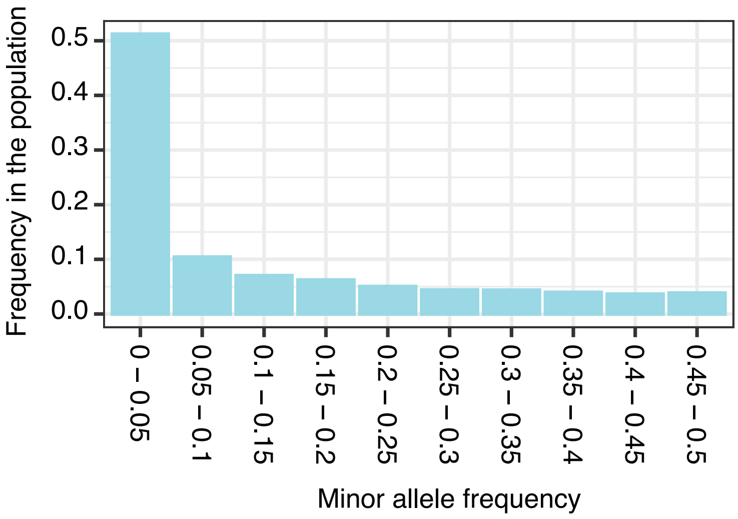


**Supplementary Figure S1.** Minor allele frequency distribution of SNPs and indels in the mapping population

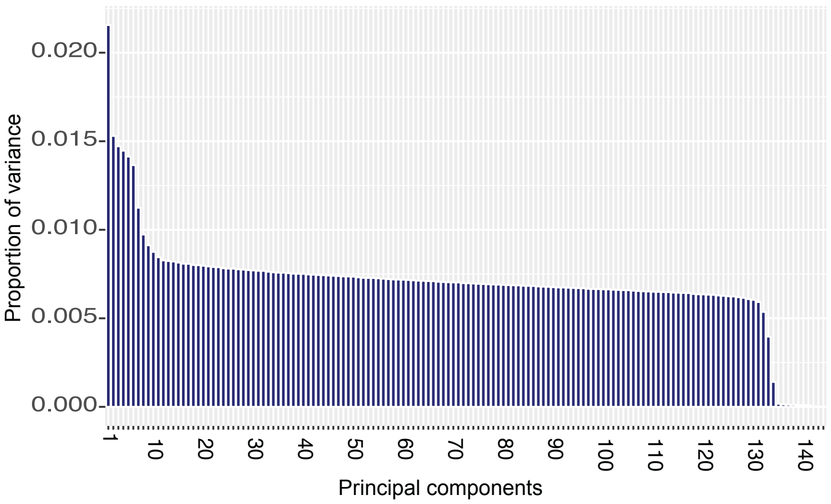


**Supplementary Figure S2.** Proportion of variance explained by the principal components based on SNP polymorphism.

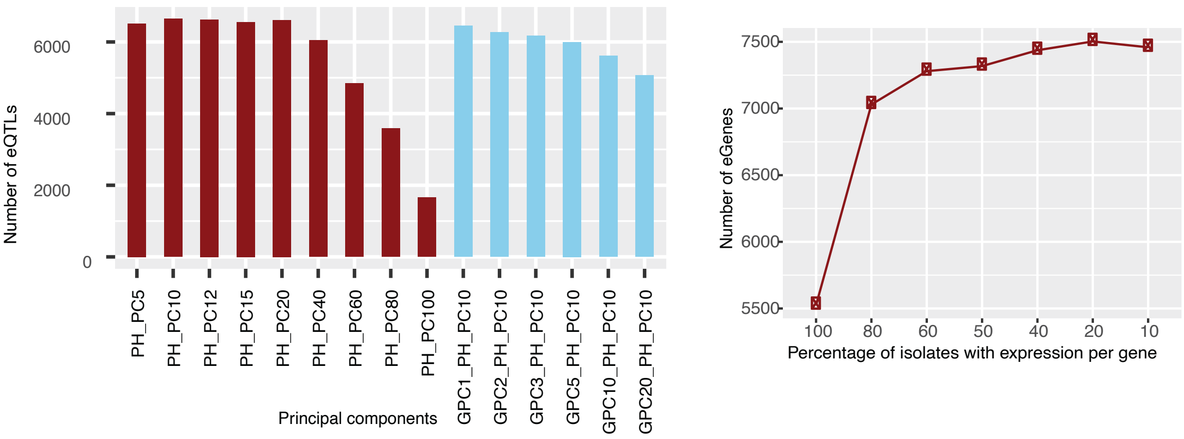


**Supplementary Figure S3. Optimization of the number of principal components to include to maximize eQTL discovery.** Left: The number of genes with eQTL reported for differing numbers of genotype (GPC) and gene expression (PH) principal components. Left: Number of genes with an eQTL mapped as a function of the required minimum percent of isolates showing gene expression.

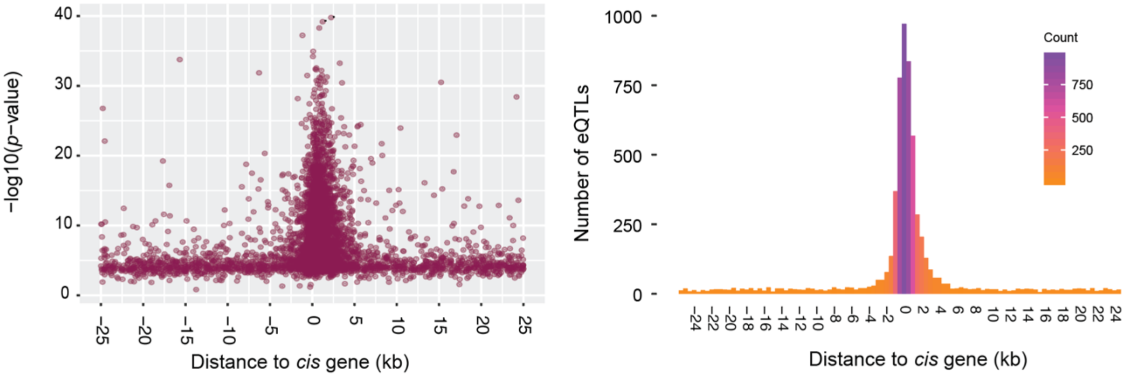


**Supplementary Figure S4. Optimization of *cis* window size around TSS for eQTL mapping.** Left: backward nominal *p*-value distribution of eQTLs mapped spanning a 25kb window centered on the TSS. Right: number of *cis*-eQTLs reported with a window size of 25kb centered on the TSS.

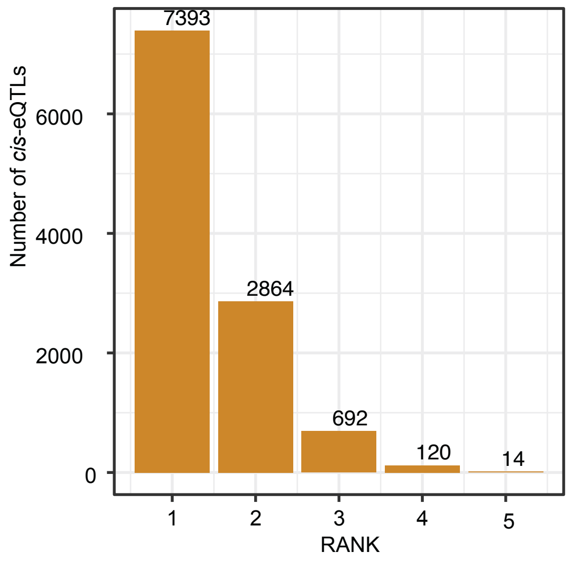


**Supplementary Figure S5.** Number of *cis*-eQTLs with decreasing effect on expression from rank 1 to rank 5.

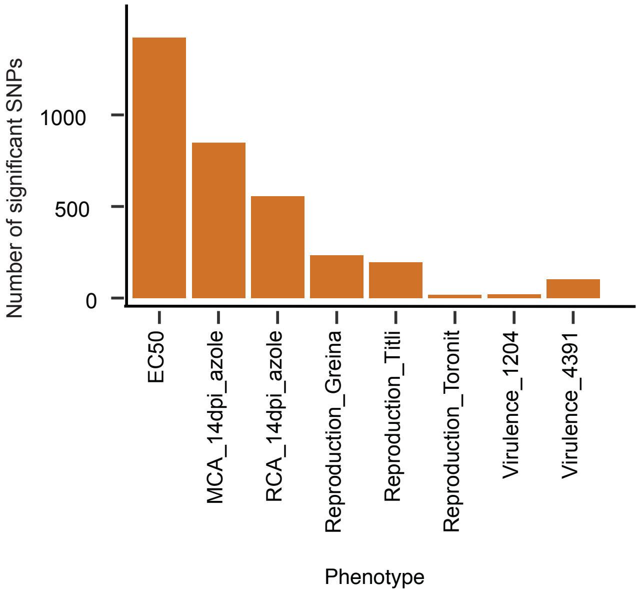


**Supplementary Figure S6.** Number of significant SNPs associated with different phenotypic traits in the mapping population.

### Supplementary Tables

**Supplementary Table S1.** Chromosomal location and statistics of all *cis*-eQTLs mapped in the mapping population.

**Supplementary Table S2.** Chromosomal location and statistics of *trans*-eQTLs identified by approximate pass.

**Supplementary Table S3**. Chromosomal location and statistics of *trans*-eQTLs identified by full pass.

**Supplementary Table S4.** Summary of virulence and fungicide sensitivity-associated genes mapped with with *trans*-eQTLs association.
